# A Facile and Versatile Technique for Creating Antifibrotic Coatings on Biomedical Implants

**DOI:** 10.64898/2026.06.04.730237

**Authors:** Yang Liu, Connor Edvall, Sudip Chakraborty, Achal Anand, Joy Agus, Suman Bose

**Author notes:** These authors contributed equally (Yang Liu, Connor Edvall).

## Abstract

Foreign body response is a common yet serious challenge for biomedical implants. It can trigger inflammation and eventually lead to the formation of a fibrotic capsule, which compromises device function. Although significant efforts have been made to develop antifibrotic surface coatings for implantable materials, developing broadly applicable solutions remains challenging due to the diversity of materials used in biomedical implants. Here, we propose a simple and versatile strategy to develop antifibrotic coatings for biomedical implants. Photoreactive benzophenone groups are incorporated into designer polymers to enable covalent attachment to various substrates. The effect of benzophenone group density within polymer chains on surface coating efficiency was investigated, and an optimal BP incorporation ratio was identified. Polymers incorporating varying ratios of an anti-fibrotic small molecule and anti-fouling zwitterionic moieties were synthesized and successfully attached to silicone implants. *In vivo* evaluation of these implants in C57BL/6 mice identified an optimized polymer composition that reduced fibrotic capsule thickness by around 60%. Coating of commercial medical catheters with this optimized polymer reduced collagen deposition by over 3.5-fold following 4 weeks of implantation in the peritoneal space of C57BL/6 mice. Finally, we demonstrated that the optimized polymer coating can be readily applied to a variety of commonly used biomedical materials using this straightforward method, highlighting the versatility of the approach. This work provides a facile and broadly applicable strategy for developing antifibrotic coatings, which has the potential to expand the design of surface modifications aimed at improving the performance of biomedical implants.

## 1. Introduction

A significant challenge for long function of biomedical implants inside the body is the foreign body response (FBR), which results in formation of a dense layer of collagen around the implant^1^. The FBR leads to a loss of implant function, such as changes in drug release kinetics^2^, instability and drift of implanted biosensors^3^, hypoxia in cell encapsulation devices^4,5^, and electrical signal interference^6^. Additionally, the fibrotic capsule can adhere to nearby tissues complicating the removal process^7^. Thus, inhibiting FBR and reducing the formation of fibrosis is necessary for the long-term function and safety of the biomedical implants.

Although hydrogels are promising biomaterials with superior biocompatibility, most medical devices used in clinics are made from plastics with suboptimal surface biocompatibility. As a result, surface modification with polymers is widely used to improve the biocompatibility of these materials and mitigate the FBR^8^. Hydrophilic polymers were among the earliest materials used to for this approach^9,10^. Due to the hydrophilic nature, a hydration layer is formed, exhibiting antifouling properties which inhibits protein adsorption and cellular attachment, preventing initiation of FBR^9^. Charged hydrophilic coatings, such as poly(acrylic acid), can also enhance antifibrotic performance through electrostatic interactions with proteins and cell membranes^11–14^. Zwitterionic polymers such as poly(2-methacryloyloxyethyl phosphocholine) (MPC), poly(sulfobetaine methacrylate) (SBMA), and poly(carboxybetaine methacrylate) (CBMA), are among the most promising antifouling materials due to their charged yet electrically neutral structures^15^. Extensive research has demonstrated the efficacy of these zwitterionic coatings in the mitigation of the FBR on various implants *in vivo*^16,17^. In recent years, high-throughput screens have been leveraged to discover several anti-fibrotic small molecules, which have demonstrated excellent performance when attached to implant surfaces in rodents and non-human primates^18,19^. Although the exact mechanisms of these anti-fibrotic small molecules remain unclear, it is likely that they modify inflammatory and tissue remodeling responses by inducing an alternative polarization of immune cells^4^. Recently, polymer coatings that combine antifouling and immunomodulatory functionalities have been shown to exhibit synergistic effects when implanted in mice, suggesting significant potential for discovering new antifibrotic materials through screening of the existing material space^20^.

The coating techniques, as well as the materials used, are equally important to develop antifibrotic coatings. Techniques using non-covalent forms of attachment, such as electrostatic interactions, hydrogen bonds, Van der Waals forces, and hydrophobic interactions, are simple but compromised with long term stability of the coating, particularly within the *in vivo* environment ^21–24^. Covalent attachment of coating polymers to implant surfaces is highly desirable, as covalent bonds offer greater resistance to degradation compared to non-covalent interactions. In the ‘graft-to’ approach, polymers are directly attached to the substrate via coupling chemistry between complementary functional groups on the polymer and substrate surface such as amine, sulfhydryl, or carboxyl groups^25^, or through click reactions such as thiol-ene or azide-alkyne cycloadditions^26^. Alternatively, in the ‘graft-from’ approach, polymers are grown directly from initiators immobilized on the substrate surface, enabling formation of dense polymer brushes^4,14^. Although a wide range of approaches exist for creating functional coatings, most current methods carry significant limitations. Coupling chemistries are highly composition dependent, requiring specific functional groups to be present on the substrate or requiring pretreatments. More commonly, coating methods require harsh reaction conditions and use of organic solvents, which may not be suitable for implantable devices carrying sensitive electronics or sensors^4^. For instance, many click chemistries use noble metal copper as the ligand which poses potential risk for *in vivo* applications^27^. When polymers are coupled to a substrate in solution, steric hindrance from already-attached polymer chains progressively limits access to the surface, constraining achievable grafting density^28^.

Stability of the attachment linker is another challenge for many of the current coating methods^29^. Silane coatings, for example, have been widely used in various coating applications^30^, however their stability can be significantly compromised under elevated temperatures and saline conditions^31,32^. This limitation poses a major challenge for long-term *in vivo* applications^33^. Therefore, there remains a need for more versatile and substrate-agnostic methods for developing antifibrotic coatings.

Incorporation of BP directly into the graft polymer provides an effective approach to integrate the versatility of BP-based photo-crosslinking with the convenience of single-step polymer coating fabrication. This method has been explored for applications in antifouling and antifog coatings, however the use of BP-based polymer for creating biocompatible coating has not been previously investigated^34–39^. In this work, we develop a potent anti-fibrotic polymer comprised of immunomodulatory and zwitterionic units along with BP acrylate (BPA) for easy attachment to a large variety of biomedical implants. We first explored the optimal BPA ratio that enhances the attachment of polymer molecules to the substrate. We found that 20% BPA incorporation into the polymer provided the best coating while retaining function. Next, we varied the ratios of MPC and THPT to create designer anti-fibrotic polymer combinations. With the creation of these polymer combinations, we demonstrated both hydrophobic and hydrophilic monomers are compatible with the technique. With the newly formed designer polymer combinations, we then assessed efficacy in vivo. We found that when 2 parts MPC, 6 parts THPT, and 2 parts BPA were combined, the resulting polymer demonstrated increased efficacy compared to the homopolymers. To confirm the efficacy and demonstrate clinical relevance, we coated ventricular catheters with the lead combination and implanted intraperitoneally. We found the lead combination resulted in an over 3.5-fold reduction in fibrosis compared to the uncoated catheter. Finally, to demonstrate the versatility of the combination, we coated four different medical substrates. We found this method can be applied to a vast variety of substrates with high density coating formation. Through this work, we aim to provide a facile and versatile option for rapid screening and development of FBR-resistant polymer coatings, with the potential of upscaling and clinical applications.

## 2. Results and Discussions

### 2.1 Developing Benzophenone based polymers and optimization of implant coating

As shown in Figure 1 (A), the developed technique enables the attachment of polymers to a variety of substrates through a UV-initiated BPA diradical coupling reaction. This method offers the flexibility of creating a wide range of polymers coatings on various substrates to further investigate their interactions with cells both in vitro and *in vivo*, providing a platform for screening novel materials with enhanced resistance to the FBR.

**Figure 1.**
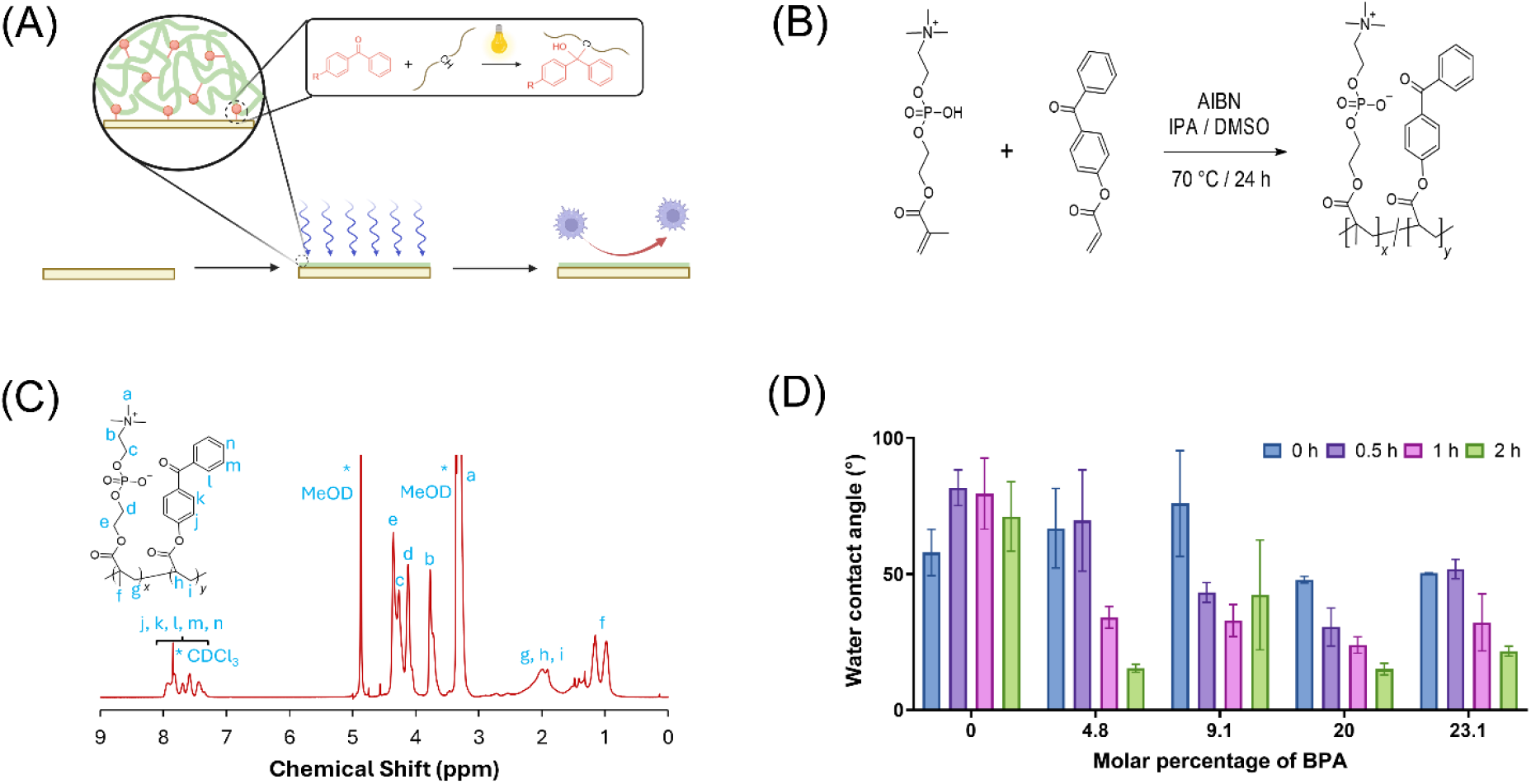
Synthesis of MPC/BPA copolymers and optimization of coating conditions. (A) The schematic of the coating process visualizes the substrate receiving the polymers, followed by attachment by UV, and finally the functionalized anti-fibrotic surface. (B) Synthetic scheme for random P(MPC-co-BPA) copolymers with the reaction conditions. (C) ^1^H NMR spectrum of a representative P(MPC-co-BPA) prepared with a feed ratio of MPC:BPA = 8:2, demonstrating successful incorporation of the BPA and MPC components. (D) Water contact angles of coatings prepared with varying BPA feed molar percentages and UV curing times.

The BPA monomer was synthesized via the esterification between acryloyl chloride and 4-hydroxybenzophenone following previous reports^34,40^and structure was confirmed by NMR (**Figure S1**). Polymers incorporating BPA attachment moieties are synthesized through conventional free-radical copolymerization, as illustrated by the representative example shown in Figure 1 (B). The corresponding ^1^H-NMR spectrum shown in Figure 1 (C) confirms the successful synthesis of the polymer. The presence of aromatic hydrogens at around 7-8 ppm indicates successful incorporation of BP monomer into the polymer. Characteristic peaks belonging to MPC, like *-C****H***_*2*_*-N-* and *-C****H***_*2*_*-O-*, appeared from 3.7 to 4.3 ppm, demonstrating the presence of MPC within the polymer. Because the BP moiety serves as the primary attachment group for polymer immobilization on the substrate, we hypothesized that both the BPA content in the polymer and the UV curing time would affect the efficiency of substrate functionalization. Therefore, the effects of the BPA ratio and UV curing time on the functionalization process were systematically evaluated. To enhance the coupling efficiency between the substrate and the BP moieties, PDMS surfaces were first modified with a layer of (3-aminopropyl)triethoxysilane (APTES) to introduce -CH-groups. Due to the hydrophilic nature of MPC, water contact angle (WCA) measurements were used to assess the extent of surface modification. Copolymers with varying MPC-to-BPA feed ratios were synthesized and immobilized onto substrates using varying UV curing times. When the molar percentage of BPA increased from 4.8% to 9.1% and then to 20.0%, the WCA of the coating surface generally decreased for a given UV curing time. However, the WCA rebounded slightly when the molar percentage of BPA increased from 20.0% to 23.1%, although both demonstrated low WCA. Surface functionalization was further confirmed by analyzing the surface elemental composition using X-ray photoelectron spectroscopy (XPS) (**Table 1**). Phosphorus is the characteristic element unique to MPC and its change is a strong indicator of the coating composition. Additionally, as the substrate uniquely contains high silicon, the decrease in this element can be used as an indicator of graft density. Thus, the success of the coatings can be confirmed by high phosphorus, while having low silicon. Consistent with WCA, the detected content of phosphorus increased with the increase of BPA when going from 4.8% to 9.1%, but it decreased when BPA increased from 9.1% to 20.0%, and further when BPA increased to 23.1%. While increasing ratios of BPA would increase the efficiency of attaching polymers onto the surface, higher ratios would decrease the effective exposure of other functional components in the copolymers. Unless specifically mentioned, 20.0 molar percent of BPA and 1 hour UV curing time are used in the subsequent experiments due to the performance and conditions of this combination being most practical.

**Table 1.** Atomic composition of P(MPC-co-BPA) coatings with varying BPA feed ratios determined by XPS.

| BPA% | C (%) | O (%) | Si (%) | N (%) | P (%) |
| --- | --- | --- | --- | --- | --- |
| Control | 26.57 ± 1.09 | 39.36 ± 1.27 | 33.67 ± 0.25 | 0.40 ± 0.08 | 0.00 |
| 4.8 | 34.30 ± 0.39 | 35.24 ± 0.38 | 27.82 ± 0.10 | 1.71 ± 0.16 | 0.92 ± 0.05 |
| 9.1 | 45.21 ± 0.45 | 30.76 ± 0.23 | 19.31 ± 0.37 | 2.99 ± 0.07 | 1.73 ± 0.16 |
| 20.0 | 44.48 ± 0.40 | 28.72 ± 0.26 | 23.92 ± 0.35 | 1.43 ± 0.21 | 1.46 ± 0.17 |
| 23.1 | 43.32 ± 1.45 | 28.35 ± 0.67 | 26.40 ± 1.72 | 1.30 ± 0.60 | 0.62 ± 0.15 |

### 2.2 Synthesis and coating of designer anti-fibrotic polymers

A key advantage to our BP-based polymers is the flexibility with which new polymers can be designed and coated on substrates. Random polymers can be designed with any number of monomer combinations to create novel coatings with diverse biological activity. After identification of the ideal BPA ratio required for substrate attachment, our next goal was to develop potent antifibrotic polymer combinations. Despite superhydrophilicity and anti-fouling properties, polyMPC (pMPC) coatings alone have failed to suppress fibrosis on biomedical devices^4^. Recent high throughput screens have discovered novel anti-fibrotic compounds with strong efficacy when used as polymer coatings^18,19^. Tetrahydropyran phenyl triazole (THPT) is a potent anti-fibrotic small molecule, and THPT modified implants have resisted fibrosis in rodent and non-human primates^4^. Recent studies have shown that combining these two classes of molecules leads to a synergistic efficacy enhancement resulting in stronger anti-fibrotic coatings^20^. Therefore, we expected that polymer combinations of MPC and THPT will lead to superior anti-fibrotic coatings compared to their homopolymer counterparts. Towards this goal, we created a range of varying MPC:THPT ratio polymers incorporating the optimized 20% BPA concentration. We synthesized and confirmed the structure of six designer anti-fibrotic polymers, with varied MPC and THPT ratios between 0 and 8 parts and 2 parts BPA. Herein, the polymers were synthesized using similar methods to the p(MPC-co-BPA) random copolymers (**Figure 2A**). NMR demonstrates the trend of signals changing with the feeding ratios (**Figure 2B**). The chemical shift at 8.18 ppm represents the triazole ring (s, *C=C****H***) that only belongs to the THPT, and chemical shifts of −*N–C****H***_*2*_*–* (2H, 3.74 ppm) and *O–C****H***_*2*_*–C****H***_*2*_*–O–P–C****H***_*2*_*–* (6H, 4.3–3.95 ppm) are characteristic peaks of pMPC.

**Figure 2.**
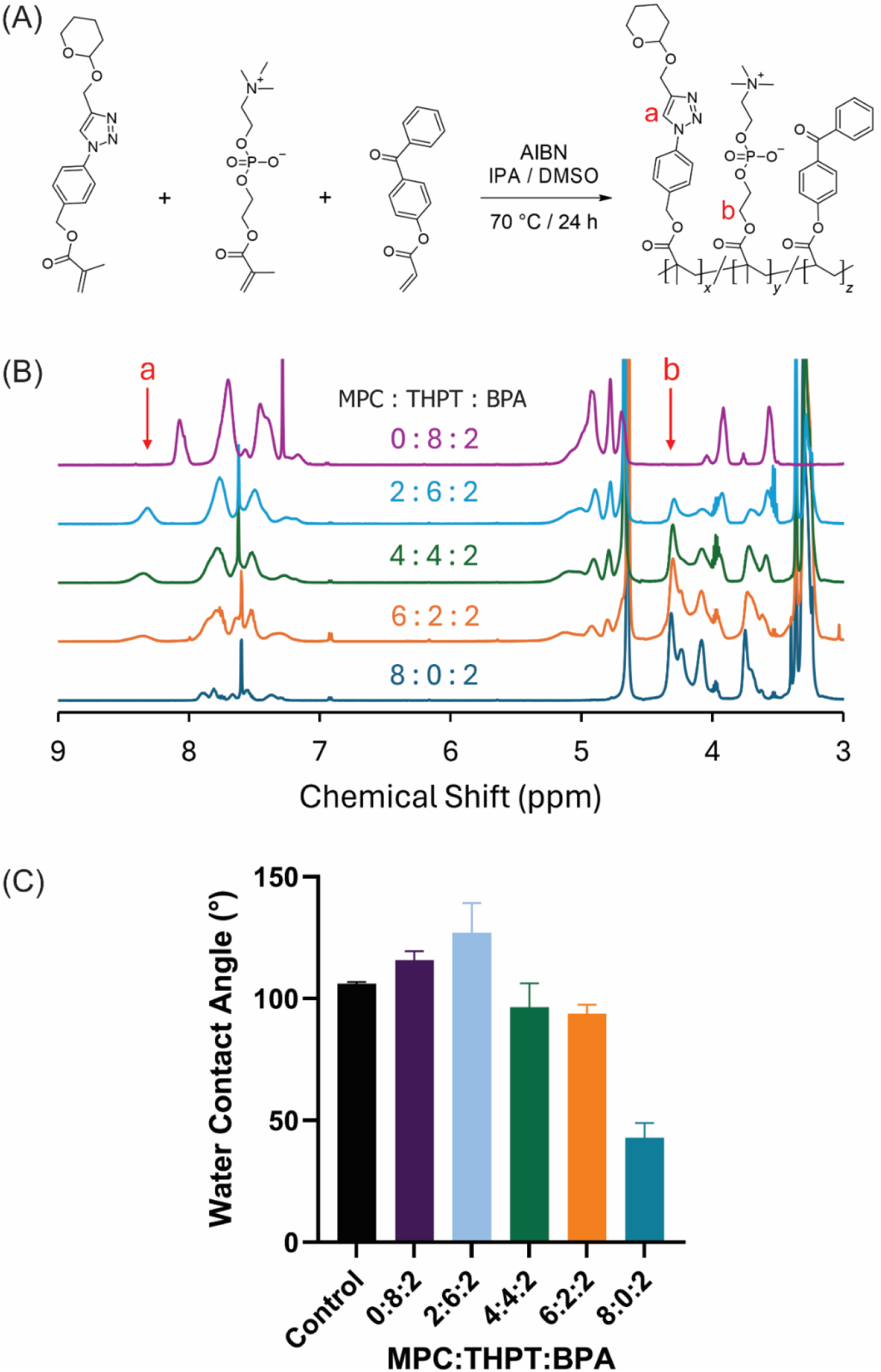
Synthesis of MPC:THPT:BPA terpolymers and coating development. (A) Synthetic scheme and reaction conditions for random P(MPC-co-THPT-co-BPA) copolymer formation. (B) ^1^H NMR spectra of copolymers prepared with varying MPC:THPT feed ratios. The characteristic THPT triazole signal at 8.18 ppm and the MPC signals at 3.7–4.3 ppm reflect changes in polymer composition with varying feed ratios. (C) Water contact angles of coatings prepared with varying monomer feed molar percentages show an overall downward trend with increasing MPC concentrations.

We observed an increasing or decreasing trend in the MPC and THPT signatures, thus confirming the composition of the copolymer could be easily controlled. While the strict control of the molecular weight of polymers is difficult via traditional radical polymerization, GPC measurements still demonstrate polymers close in weight (**Figure S4**). The synthesized copolymers were then coated on 5mm PDMS discs for further WCA and XPS analysis. Figure 2(C) shows that increasing the MPC content leads to lower WCA, with the MPC/BPA copolymer exhibiting a WCA as low as 40°. In contrast, increasing the THPT content results in higher WCA, with the THPT copolymer reaching approximately 115°, exceeding that of the substrate. This trend is consistent with the expected effect of composition on surface wettability, as zwitterionic MPC promotes water affinity, whereas THPT, containing extensive aromatic and aliphatic segments, reduces it. Interestingly, the polymer coating with feeding ratio of 2:6:2 (MPC:THPT:BPA) has the largest WCA implying unexpected interactions between different polymer components. The successful coatings were further confirmed by XPS and visualized by elemental percentage (**Table 2**). The atomic percentage of phosphorous increased from 0.45 ± 0.07% to 1.46 ± 0.07% when the feeding ratio of MPC in the polymer increased from 20% to 80% (**Table 2**). We further performed an elemental scan for N1s to determine the chemical structure of nitrogen which shows the signature of quaternary amine N^+^(CH_3_)_3_ at 402.8 eV, representing MPC, and the triazole ring from THPT at 400.4 (<u>N</u>-N=N) and 401.9 Ev (N-<u>N</u>=N and N-N=<u>N</u>) (**Figure S6**).

**Table 2.** Surface elemental composition of coatings prepared from copolymers synthesized using different monomer feed ratios, as determined by XPS.

| MPC:THPT:BPA | C (%) | O (%) | Si (%) | N (%) | P (%) |
| --- | --- | --- | --- | --- | --- |
| Control | 36.67 ± 0.60 | 30.34 ± 0.23 | 32.54 ± 0.30 | 0.45 ± 0.09 | 0.00 |
| 0 : 8 : 2 | 53.40 ± 0.97 | 21.89 ± 0.55 | 22.52 ± 0.60 | 2.18 ± 0.17 | 0.00 |
| 2 : 6 : 2 | 56.10 ± 0.36 | 21.02 ± 0.12 | 19.94 ± 0.49 | 2.48 ± 0.17 | 0.45 ± 0.07 |
| 4 : 4 : 2 | 60.66 ± 0.52 | 21.04 ± 0.27 | 13.70 ± 0.41 | 3.24 ± 0.11 | 1.36 ± 0.05 |
| 6 : 2 : 2 | 52.29 ± 3.39 | 24.82 ± 1.11 | 19.38 ± 2.73 | 2.14 ± 0.27 | 1.37 ± 0.16 |
| 8 : 0 : 2 | 44.48 ± 0.16 | 28.72 ± 0.11 | 23.92 ± 0.14 | 1.43 ± 0.09 | 1.46 ± 0.07 |

The quaternary amine was only observed when THPT is absent from the polymer, while the triazole group only appears when MPC is absent. Therefore, both peaks are integrated when terpolymers are coated, and the relative intensity of peak follows their contents. The quantitative element analysis and the qualitative high-resolution spectra (**Figure S6**) demonstrate that polymers with different compositions were successfully coated on the surface.

### 2.3 In vivo screening of copolymer coating libraries uncovers optimal anti-fouling and immunomodulatory balance for implant biocompatibility

The polymer library synthesized from THPT and MPC monomers was coated onto 5 mm PDMS discs and implanted subcutaneously in C57BL/6 mice. To reduce animal usage, we employed a high-throughput implantation strategy adapted from prior work^18^, in which each animal receives multiple implants simultaneously (Figure 3a). Implants were harvested at 4 weeks, the established timepoint for stable fibrotic capsule formation in rodents. Explanted devices and surrounding tissue were processed for histological analysis, and fibrotic capsule thickness was quantified from Masson’s Trichrome-stained sections at both the skin-facing and muscle-facing interfaces.

**Figure 3.**
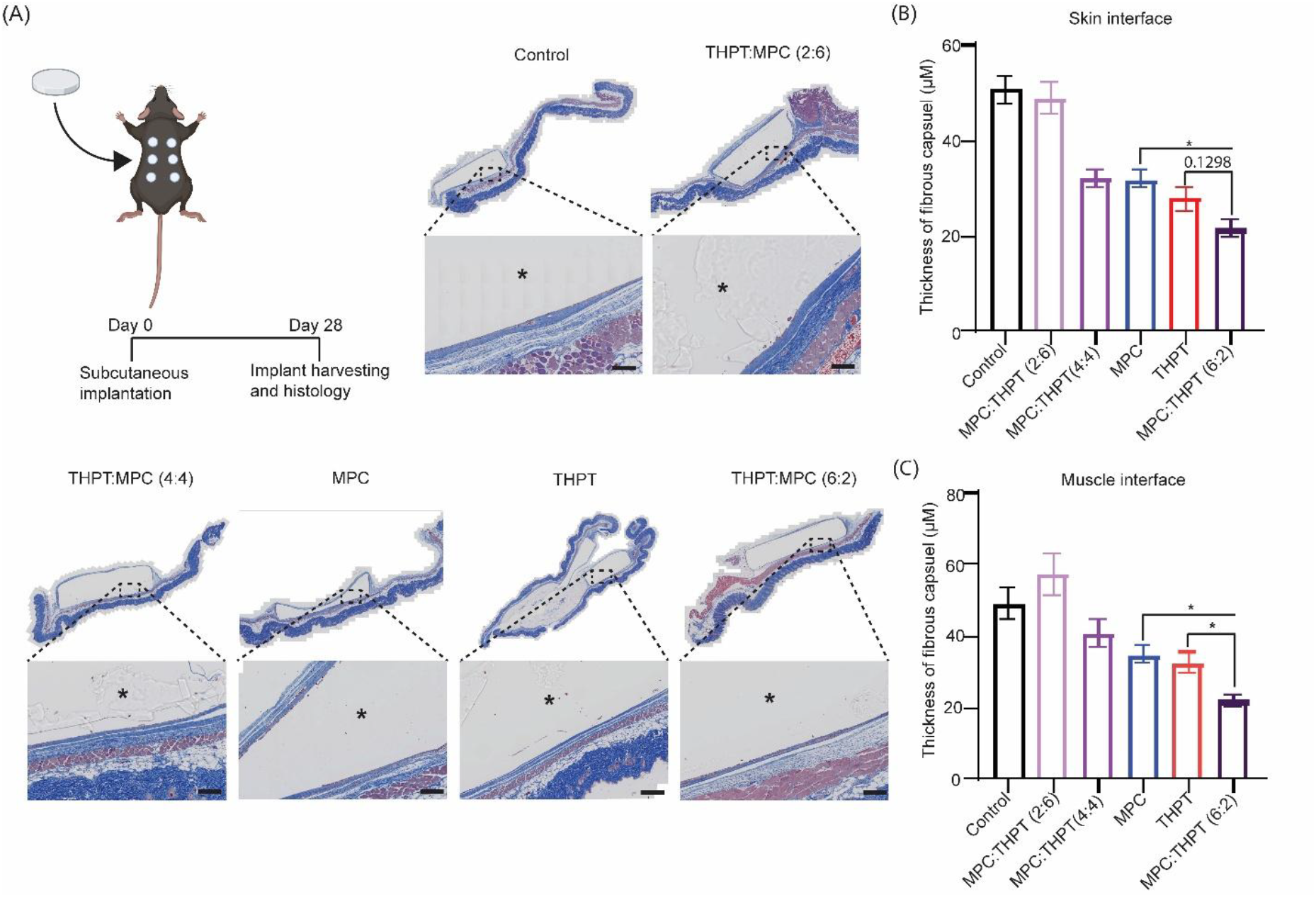
The antifibrotic performances of the MPC:THPT:BPA coatings on PDMS disks in C57BL/6 mice. (**a**) Timeline and schematic of the *in vivo* fibrosis assessment and representative histology images of the disks post-harvest, with the full image on top and the zoomed in section used for quantification on the bottom. (**b**) Quantified fibrotic capsule thickness on the skin facing interface after harvest demonstrates the efficacy of the coatings. Bar heights and error bars represent mean ± s.e.m, n = 6 implants per group. (**c**) Quantified fibrotic capsule thickness on the muscle facing interface after harvest shows a greater difference in coating performance of the hit formulation compared to homopolymers. Bar heights and error bars represent mean ± s.e.m, n = 6 implants per group (*P<0.05, One Way ANOVA). Scalebar = 100 micron.

MPC and THPT homopolymer coatings each reduced fibrotic capsule thickness by approximately 40% relative to uncoated controls. Strikingly, the MPC:THPT (6:2) copolymer significantly outperformed either component alone, achieving around a 60% reduction in capsule thickness. All other copolymer ratios performed comparably to or worse than the individual homopolymers. These results compare favorably to a previously reported block copolymer architecture combining MPC and THPT on PDMS, which achieved ∼45% reduction and an absolute capsule thickness of ∼30 µm^20^. Our optimized formulation exceeded this benchmark, yielding ∼60% reduction and a mean capsule thickness of ∼20 µm. This is likely due to the difference in polymer architecture and high density of polymer grafting density as supported by XPS measurements.

To validate the lead formulation, ventricular catheters (barium impregnated silicon) were cut into 10 mm segments, coated with the lead (2:6:2 MPC:THPT:BPA) formulation, and implanted intraperitoneally in C57BL/6 mice. After 4 weeks, the catheters were harvested, processed for histology, and stained with H&E and Masson’s Trichrome (**Figure 4B**). Quantification of the capsule thickness demonstrates an over 3.5 times reduction in the capsule thickness when coated with the lead formulation compared to the control (**Figure 4C**). Overall, these results establish the effectiveness of the coating technique *in vivo*, the ability to ‘plug and play’ with various formulations, and the applicability of the method to medical substrates.

**Figure 4.**
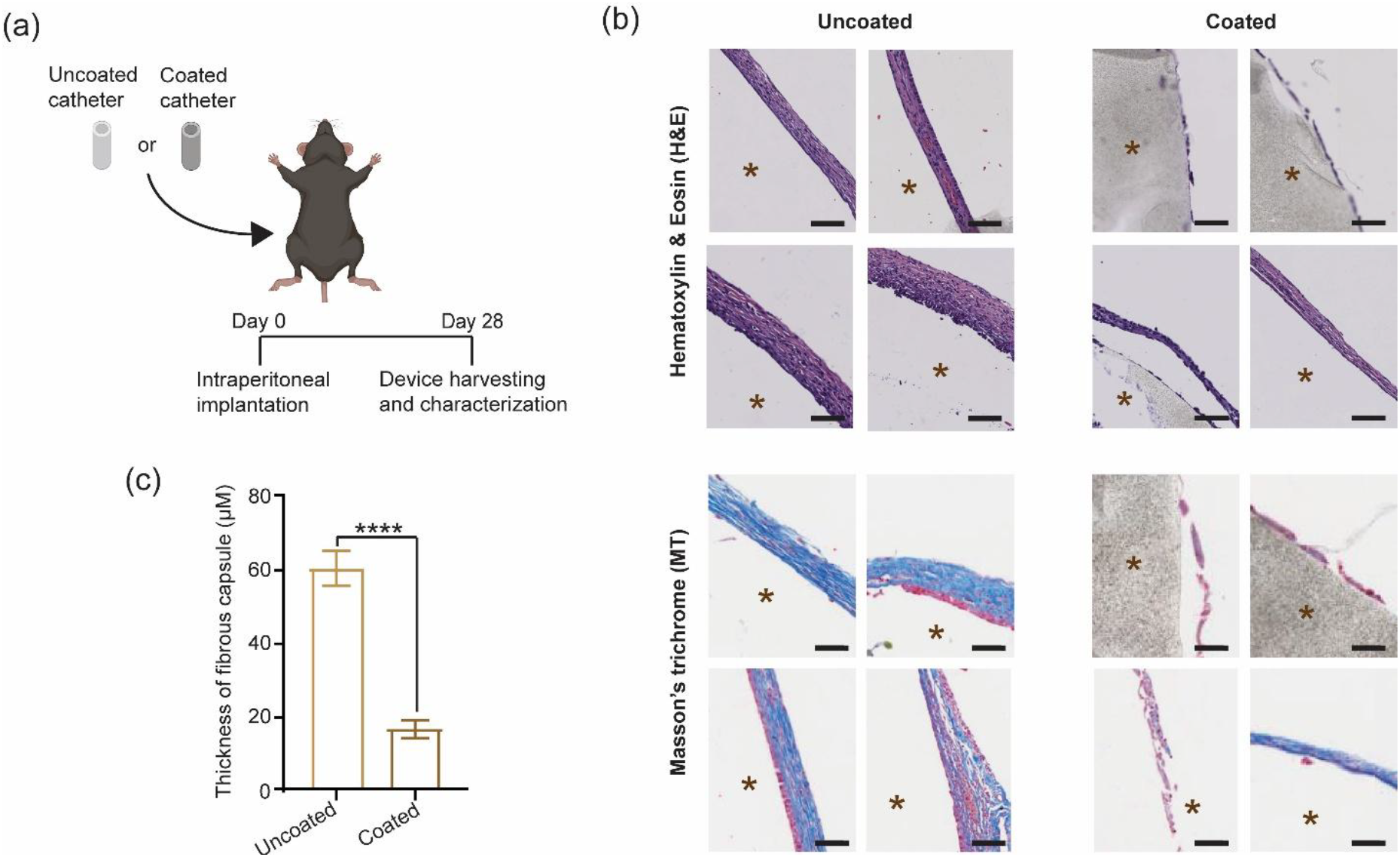
The antifibrotic performance of the 2:6:2 MPC:THPT:BPA coating on ventricular catheters in C57BL/6 mice. (**a**) Timeline of the catheter *in vivo* assessment. (**b**) Histology images of the catheters post-harvest were stained with H&E or MT. (**c**) Quantified fibrotic capsule thickness on the catheters post-harvest demonstrates the efficacy of the coating over the control. Bar heights and error bars represent mean ± s.e.m, n = 4 implants per group (****P<0.0001, One Way ANOVA). Scalebar = 50 micron.

### 2.4 Versatility of the coating technique

A key advantage of our coating technique is the versatility of substrates in which it is applicable. Because of the BP chemistry being able to form bonds with multiple substrates, we hypothesized it could be applied to a diverse range of biomedical materials. To assess this, we coated four substrates which are commonly used for medical applications and confirmed successful attachment with XPS. We found the coating technique resulted in successful coating of every substrate tested. The versatility of the coating technique was evaluated on polyurethane (PU), polymethyl methacrylate (PMMA), ventricular catheters, and polycarbonate (PC). PU is used for a variety of biomedical applications, including tissue engineering, artificial organs, surgical sutures, and medical catheters^41^, PMMA is commonly used as a bone cement in orthopedic surgery^42,43^, ventricular catheters are used to primarily for diversion of cerebrospinal fluid^44^, and PC is frequently used in implantable device components^1^.To evaluate the coating success, the surface elemental compositions of PU, PMMA, PC, and ventricular catheters were analyzed before and after coating with the lead formulation (2:6:2 MPC:THPT:BPA) using XPS (**Table 3**). The nitrogen and phosphorus content were used as characteristic elements of MPC and THPT due to either low or no content in the substrates. We observed a substantial increase in both elements, confirming a dense coating on multiple substrates.

**Table 3.** Atomic composition of the lead coating applied to various biomedical substrates.

|  | C (%) | O (%) | Si (%) | N (%) | P (%) |
| --- | --- | --- | --- | --- | --- |
| Ventricular Catheter | 51.10 ± 0.19 | 21.61 ± 0.04 | 26.12 ± 0.20 | 1.18 ± 0.04 | 0.00 |
| Ventricular Catheter-Coating | 60.45 ± 0.19 | 20.10 ± 0.10 | 15.49 ± 0.16 | 3.55 ± 0.11 | 0.41 ± 0.03 |
| PU | 83.11 ± 0.09 | 14.10 ± 0.07 | 0.00 | 2.80 ± 0.12 | 0.00 |
| PU-Coating | 74.35 ± 0.07 | 19.21 ± 0.15 | 0.00 | 5.24 ± 0.19 | 1.20 ± 0.12 |
| PMMA | 76.76 ± 0.14 | 23.24 ± 0.14 | 0.00 | 0.00 | 0.00 |
| PMMA-Coating | 74.78 ± 0.05 | 17.08 ± 0.06 | 0.00 | 6.66 ± 0.06 | 1.49 ± 0.02 |
| PC | 85.53 ± 0.11 | 14.47 ± 0.11 | 0.00 | 0.00 | 0.00 |
| PC-Coating | 74.56 ± 0.10 | 17.28 ± 0.05 | 0.00 | 6.72 ± 0.06 | 1.43 ± 0.01 |

Our results indicate the coating method is applicable on a variety of common biomedical materials, despite differences in their chemical structures. Thus, demonstrating the versatility of this coating strategy and highlighting its potential for developing antifibrotic coatings for a wide range of implantable medical devices.

## 3. Conclusions

In this work, we developed a simple and versatile strategy for constructing antifibrotic coatings on biomedical implants. We first established the optimal BPA to MPC ratio, allowing maximum coating efficiency while retaining efficacy. We then created designer antifibrotic polymers and screened them *in vivo* to identify the lead formulation. This formulation then demonstrated around a 60% decrease in fibrosis on PDMS, and an over 3.5-fold reduction on ventricular catheters. Finally, we demonstrated the versatility of the method by coating four biomedical substrates with a dense polymer layer.

Overall, our work demonstrates a facile and versatile anti-fibrotic coating strategy for biomedical substrates. Coatings could be applied to materials which are prone to loss of function, such as biosensors^45,46^, various orthopedic or cosmetic implants^47,48^, and immune isolation devices^4^ for future applications. Additionally, this technique can provide additional advantages over current strategies allowing a gateway for commercial translation. It is clean room compatible, reduces coating times from hours to minutes, applicable to a variety of substrates and solvents, and is readily scalable for large scale production, demonstrating both clinical and commercial potential.

## 4. Materials and Methods

### 4.1 Materials

4-Hydroxybenzophenone, acryloyl chloride, triethylamine, 2-methacryloyloxyethyl phosphorylcholine, azobisisobutyronitrile (AIBN), (3-aminopropyl)triethoxysilane (APTES), methanol, ethanol, isopropanol, 2-butanone, tetrahydrofuran (THF), dichloromethane (DCM), dimethyl sulfoxide (DMSO), sodium hydroxide (NaOH), and sodium sulfate (Na_2_SO_4_) were purchased from Sigma-Aldrich and used without further purification. Tetrahydropyran phenyl triazole (THPT) monomers were synthesized by WuXi AppTec Co., Ltd.

PDMS substrates were prepared by mixing SYLGARD™ 184 silicone elastomer base and curing agent (Dow Chemical Company) at a mass ratio of 10:1 and curing at 60 °C for 24 h. Silicone catheters (CODMAN HOLTER Atrial Catheter Type A) were provided by Integra LifeSciences Production Corporation. Polycarbonate (PC) sheets were purchased from Goodfellow Advanced Materials. Single-lumen polyurethane (PU) catheters were purchased from VYGON Value Life. Poly(methyl methacrylate) (PMMA) sheets were provided by Astra Products Inc.. BPA monomers were synthesized following previously reported procedures^34,40^, and the corresponding NMR spectra are provided in the Supporting Information (Figure S1).

### 4.2 Synthesis of polymers

Random copolymers and terpolymers were synthesized via radical polymerization adapted from previous research. Monomers and the initiator AIBN, at the designed ratios, were dissolved in a cosolvent mixture of isopropanol and DMSO (7:3). Oxygen in the solution was removed by three freeze–pump–thaw cycles prior to polymerization at 70 °C for 24 h. The obtained polymers were precipitated twice in diethyl ether and then dried under vacuum for 24 h.

The polymers were characterized by NMR. Polymers containing MPC were dissolved in a cosolvent mixture of methanol-d_4_ and chloroform-d, whereas polymers without MPC were dissolved in pure chloroform-d. ^1^H NMR spectra were recorded on a Bruker Avance NEO 500 MHz spectrometer with 32 scans. The molecular weights of the polymers were measured using an Agilent 1260 Infinity II system with a methanol/water (1:1) cosolvent.

### 4.3 Development of surface coatings

For surfaces only containing methyl groups (e.g., PDMS or silicone), an APTES silane layer was introduced to improve coating attachment. Substrates were treated with oxygen plasma for 60 s and immediately immersed overnight in a solution of APTES (2% v/v), water (3% v/v), and ethanol (95% v/v). The substrates were then thoroughly rinsed with ethanol and dried at 60 °C for 1 h. The silane-pretreated substrates were used for subsequent coating experiments. The solvent and polymer concentration were adjusted based on the compatibility among the polymer, solvent, and substrate. MPC–BPA copolymers were dissolved in ethanol (15 mg/mL) for coating on PDMS. Copolymers and terpolymers of MPC, THPT, and BPA were dissolved in THF/methanol (1:1) to obtain 15 mg/mL coating solutions for silicone catheters. A terpolymer of MPC, THPT, and BPA with a feed ratio of 2:6:2 was dissolved in butanol/methanol (1:1) to obtain a 4 mg/mL solution for coating on PC, PMMA, and PU. Prior to coating, all substrates were treated with oxygen plasma for 15 s to increase surface wettability. The plasma-treated substrates were immediately immersed in the polymer solutions, removed, and dried under ambient conditions. This process was repeated twice to improve coating coverage. Coatings on flat substrates were cured under 365 nm UV irradiation (15 mW/cm^2^) for 1 h. Coatings on cylindrical catheters were cured for 40 min, rotated by 90°, and cured for an additional 40 min; this step was repeated once more to ensure complete surface coverage. The resulting coatings were washed with ethanol for 1 h, followed by PBS overnight, then thoroughly rinsed with deionized water and dried under a nitrogen stream.

Water contact angles were measured to characterize the hydrophilicity of the coatings. A Theta Lite Optical Tensiometer (Biolin Scientific) equipped with OneAttension software was used to measure the water contact angles of the coatings. A 5 µL droplet of deionized water was placed on the surface for imaging, and each sample was measured at three different locations to obtain the mean value.

Surface elemental composition was analyzed using an X-ray photoelectron spectrometer (Kratos Axis Supra+) equipped with a monochromatic Al Kα X-ray source. The binding energy (BE) range of the survey spectrum was from 1300 to 0 eV, with a pass energy of 80 eV. The obtained spectra were analyzed using CasaXPS (Casa Software Ltd.).

### 4.4 In-vivo Experiments

All animal procedures were approved by Mayo Clinic Institutional Animal Care and Use Committee prior to starting. Male C57BL/6 mice of 6-8 weeks in age were obtained from The Jackson Laboratory to be used in the study. Prior to surgery, mice were anaesthetized with 2.5% isoflurane in oxygen. Subsequently, mice had either their back or abdomens shaved and sterilized via betadine and isopropyl alcohol. 3.25 mg/kg Buprenorphine (Ethiqa XR) and 1 mL 0.9% saline were injected subcutaneously. For the subcutaneous surgeries, six small incisions were made with three on each side of the back. PDMS disks were then inserted into the incisions, and the wounds closed using wound clips. For the intraperitoneal surgery, a small incision was made along the midline of the abdomen. An incision was then made alone the linea alba. The catheters were then inserted into the abdomen and orientated away from the fat pad. The peritoneum was then closed using a 6-0 size absrobable vicryl suture. The incision site was then closed using wound clips.

Retrieved implants were fixed for 20 min using 4% paraformaldehyde at room temperature. Implants were then washed twice in PBS, cut in half, then stored in 70% ethanol. The blocks were processed for paraffin embedding with care to ensure the samples were orientated for even cross sectioning of the implants. The blocks were sectioned and stained using standard histology techniques. The fibrotic capsule thickness was quantified on the skin and muscle interface using at least 5 sites per side with QuPath (version 0.6.0) software. The MPC and THPT controls were then compared to the best ratio using one way ANOVA.

## Acknowledgements

Research reported in this publication was supported by the National Institute of General Medical Sciences of the National Institutes of Health under award number R35GM155031 and Mayo Clinic Office of Core Shared Services Early-Stage Investigator Research Award to S.B. The authors also acknowledge resources and support from the Magnetic Resonance Research Center, part of the Chemical and Environmental Characterization Core Facilities at Arizona State University, and the Eyring Materials Center at Arizona State University supported in part by NNCI-ECCS-1542160. We would also like to thank Thang Phan for assisting with the synthesis.

## Ethics Approval Statement

All animal procedures were approved by Mayo Clinic Institutional Animal Care and Use Committee prior to starting.

## Data Availability Statement

The data that supports the findings presented in this study are available from the corresponding author upon request.

