## Supplementary Information for "A Facile and Versatile Technique for Creating Antifibrotic Coatings on Biomedical Implants"

[+]These authors contributed equally

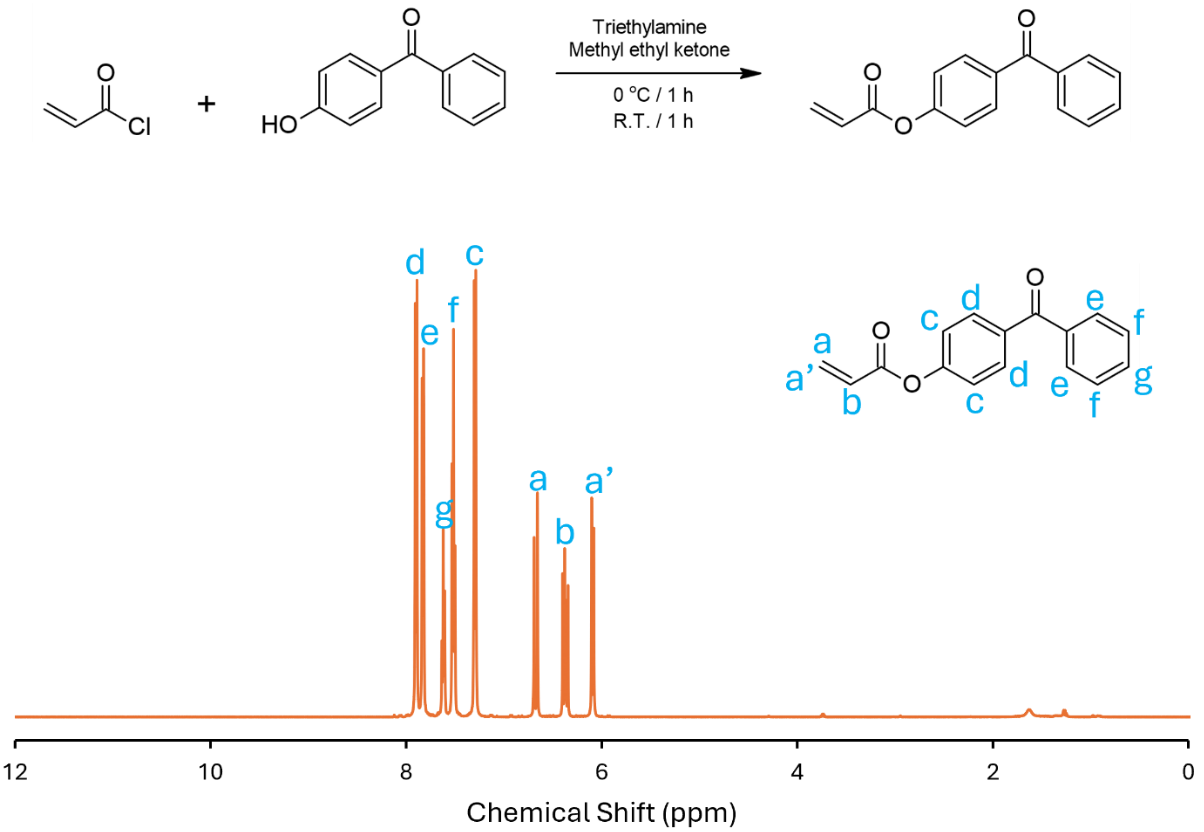


**Figure S1**. Synthesis scheme of BPA and its corresponding $^{1}$H NMR spectrum.

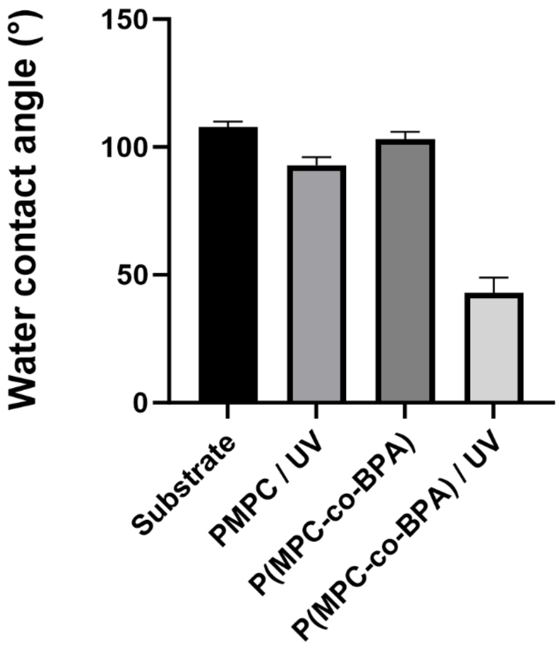


**Figure S2**. Water contact angles of polymer coatings in the presence or absence of BPA and UV Curing.

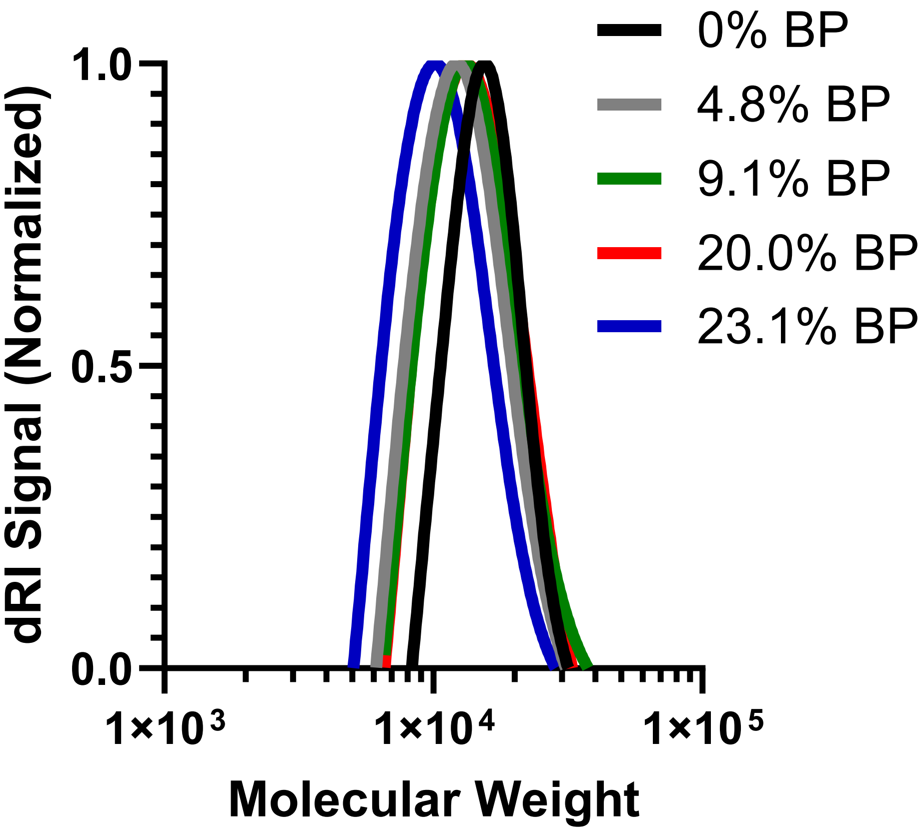


**Figure S3**. Distribution of varied MPC:BP polymer combination molecular weights confirmed via GPC.

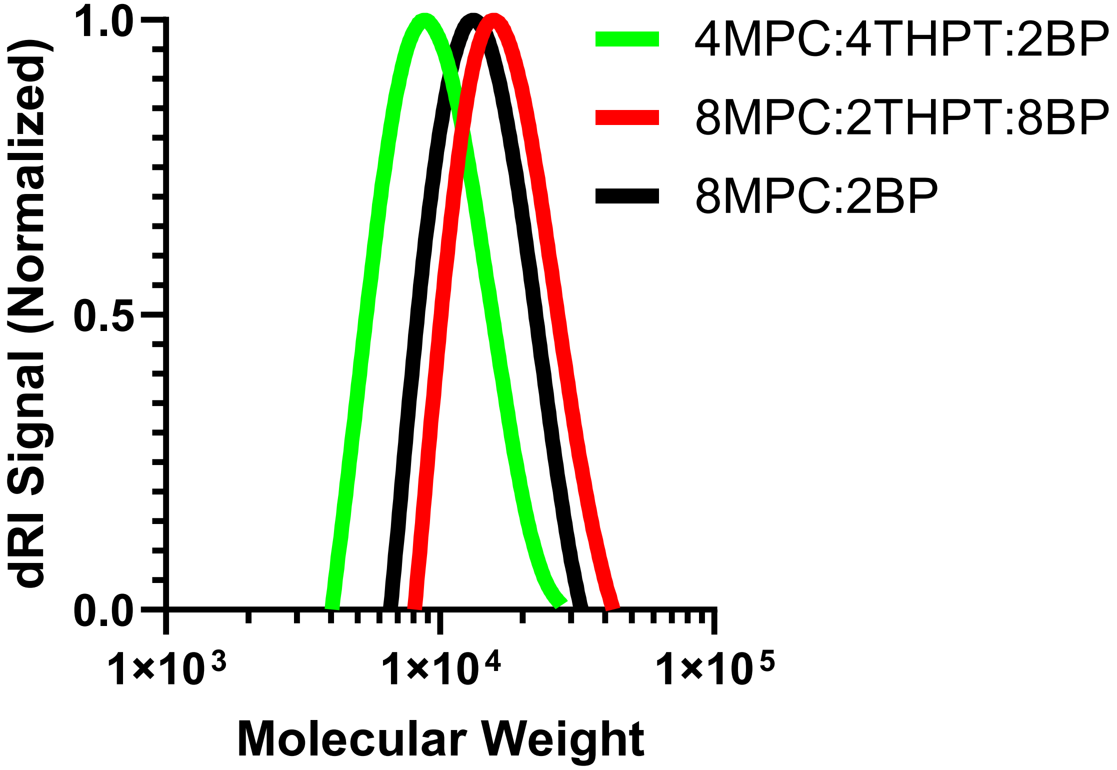


**Figure S4**. Distribution of varied MPC:THPT:BP polymer combination molecular weights confirmed via GPC.

**Table S1**. Molecular weight characteristics of copolymers prepared with different feed ratios. * indicates incompatibility with aqueous and organic phase GPC.

| MPC:THPT:BP | 0 : 8 : 2[a] | 2 : 6 : 2[a] | 4 : 4 : 2[b] | 6 : 2 : 2[b] | 8 : 0 : 2[b] |
|---|---|---|---|---|---|
| $M_n$ | * | * | 13600 | 15700 | 13100 |

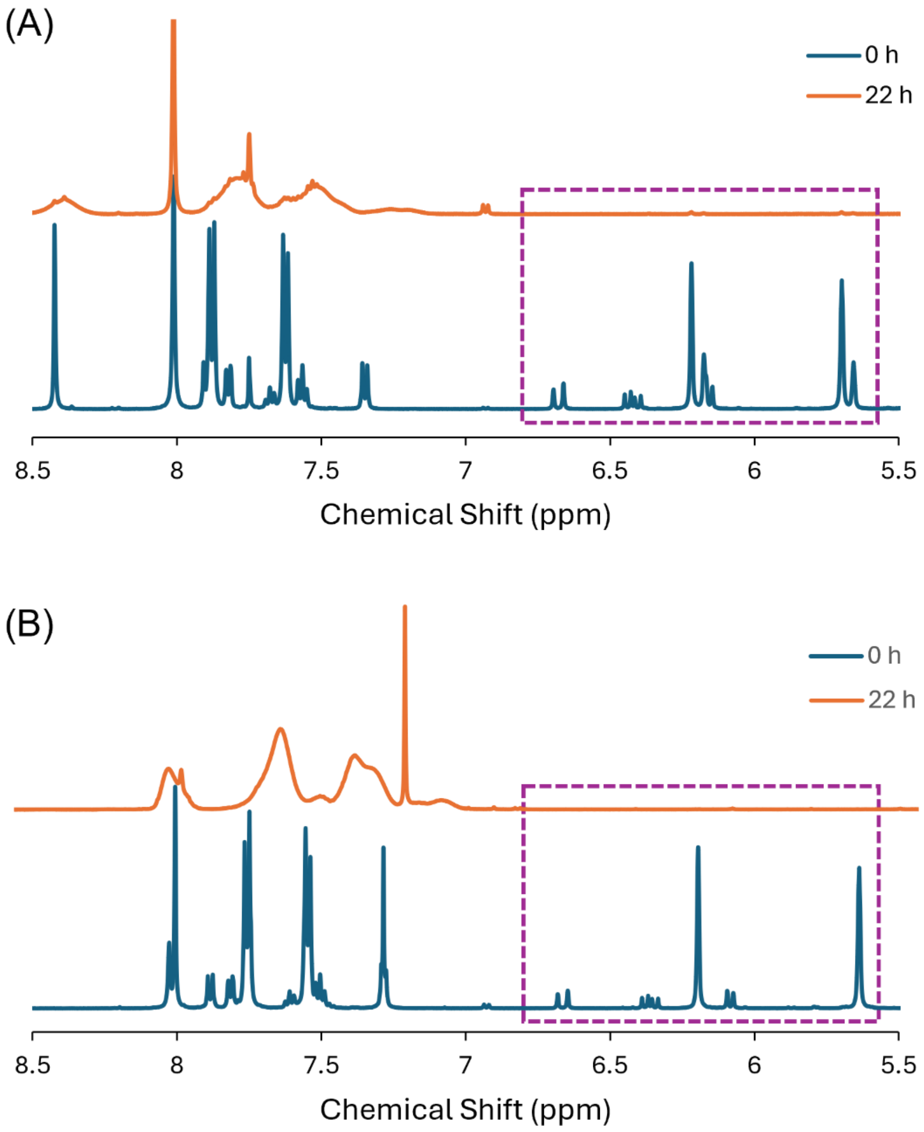


**Figure S5**. Monomer consumption of synthesis MPC : THPT : BPO = 2 : 6 : 2 (A) and MPC : THPT : BPO = 0 : 8 : 2 (B). The disappearance of double bond within 5.5 – 7.0 ppm indicating the consumption of monomers.

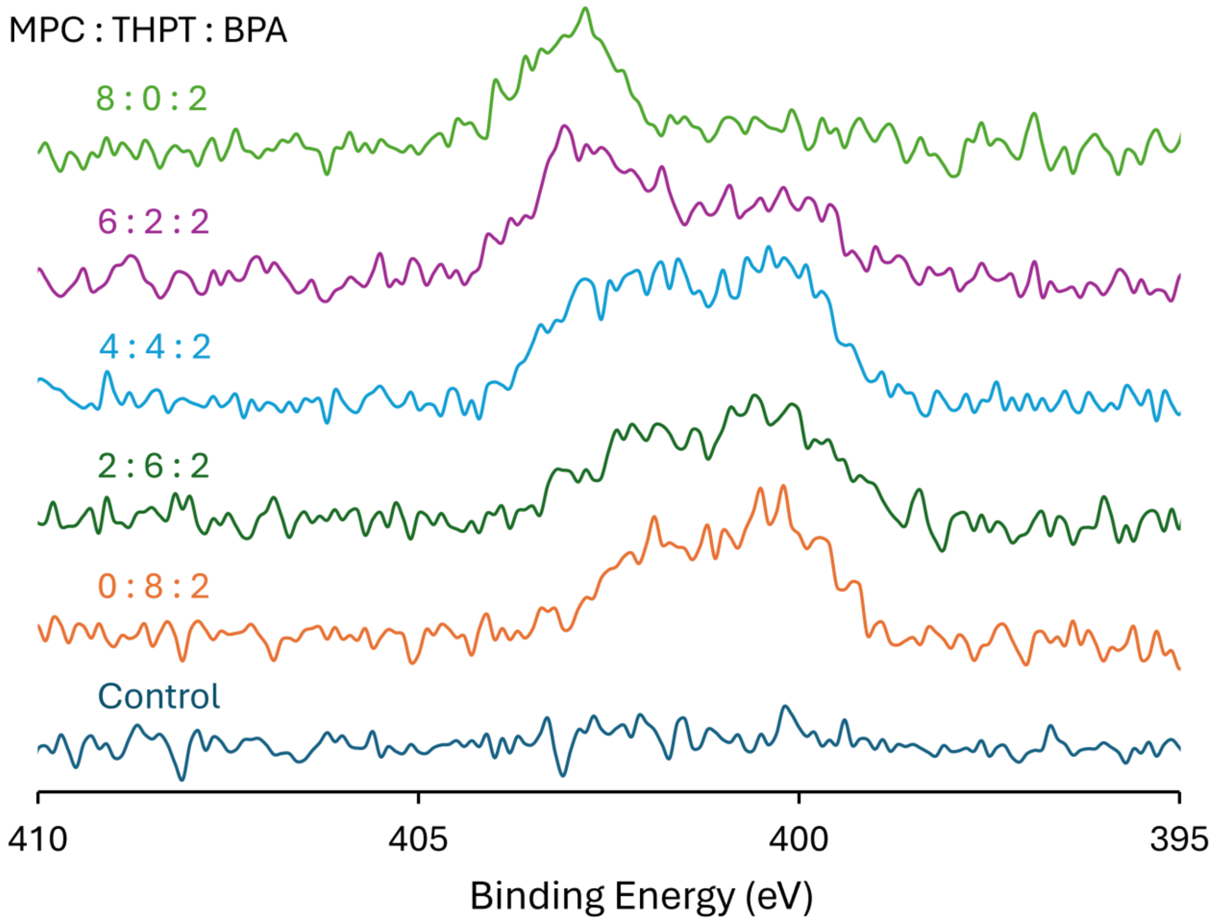


**Figure S6**. High resolution XPS spectra of polymer coatings prepared with different feed ratios.
